# Junctophilin-2 Regulates Mitochondrial Metabolism

**DOI:** 10.1101/2023.02.07.527576

**Authors:** Sasha Z. Prisco, Lynn M. Hartweck, Felipe Kazmirczak, Jenna B. Mendelson, Stephanie L. Deng, Satadru K. Lahiri, Xander H.T. Wehrens, Kurt W. Prins

## Abstract

Right ventricular dysfunction (RVD) is a risk factor for mortality in multiple cardiovascular diseases, but approaches to combat RVD are lacking. Therapies used for left heart failure are largely ineffective in RVD, and thus the identification of molecules that augment RV function could improve outcomes in a wide-array of cardiac limitations. Junctophilin-2 (JPH2) is an essential protein that plays important roles in cardiomyocytes, including calcium handling/maintenance of t-tubule structure and gene transcription. Additionally, JPH2 may regulate mitochondrial function as *Jph2* knockout mice exhibit cardiomyocyte mitochondrial swelling and cristae derangements. Moreover, JPH2 knockdown in embryonic stem cell-derived cardiomyocytes induces downregulation of the mitochondrial protein mitofusin-2 (MFN2), which disrupts mitochondrial cristae structure and transmembrane potential. Impaired mitochondrial metabolism drives RVD, and here we evaluated the mitochondrial role of JPH2. We showed JPH2 directly interacts with MFN2, ablation of JPH2 suppresses mitochondrial biogenesis, oxidative capacity, and impairs lipid handling in iPSC-CM. Gene therapy with AAV9-JPH2 corrects RV mitochondrial morphological defects, mitochondrial fatty acid metabolism enzyme regulation, and restores the RV lipidomic signature in the monocrotaline rat model of RVD. Finally, AAV-JPH2 improves RV function without altering PAH severity, showing JPH2 provides an inotropic effect to the dysfunction RV.

---

Right ventricular dysfunction (RVD) is a risk factor for mortality in multiple cardiovascular diseases, but approaches to combat RVD are lacking^1^. Therapies used for left heart failure are largely ineffective in RVD, and thus the identification of molecules that augment RV function could improve outcomes in a wide-array of cardiac limitations. Junctophilin-2 (JPH2) is an essential protein that plays important roles in cardiomyocytes, including calcium handling/maintenance of t-tubule structure and gene transcription^2^. Additionally, JPH2 may regulate mitochondrial function as *Jph2* knockout mice exhibit cardiomyocyte mitochondrial swelling and cristae derangements^3^. Moreover, JPH2 knockdown in embryonic stem cell-derived cardiomyocytes induces downregulation of the mitochondrial protein mitofusin-2 (MFN2), which disrupts mitochondrial cristae structure and transmembrane potential^4^. Impaired mitochondrial metabolism drives RVD^1^, but JPH2’s potential interaction with MFN2 and its impact on RV mitochondrial activity are not well-defined.

Cellular fractionation, co-immunoprecipitation, and pulldown assays of recombinantly expressed and purified JPH2 and MFN2 constructs defined a direct JPH2-mitochondrial link. *E. coli*-codon optimized JPH2 was synthesized and cloned into pET151/dTOPO (Thermo) with deletions generated by Q5 mutagenesis PCR (NEB). His-tagged proteins were purified and tested for interaction by co-immunoprecipitation with MFN2 (Abcam), JPH2 (Invitrogen), isotype IgG, and V5 (VWR) antibodies. Human induced pluripotent stem cells (iPSC) (Allen Institute) were treated with CRISPR-Cas9 to knockout *JPH2* (Synthego). Cell proliferation, nucleofection, and differentiation of iPSC into cardiomyocytes (iPSC-CM) was performed as recommended by the Allen Institute. Super resolution microscopy [Tom20 antibody (Abcam)] defined mitochondrial density in iPSC-CM. Agilent XFp Seahorse Mito Stress examined mitochondrial respiratory (3 μM oligomycin, 20 μM FCCP, 1 μM antimycin A, and 1 μM rotenone). *In vitro* lipid sensitivity was probed by incubating iPSC-CM with 16 μM oleate, linoleate, and palmitate (Sigma) overnight. Lipid droplets were visualized with LipidTOX Red (FisherScientific). Confocal micrographs were collected on a Zeiss LSM 900 Airyscan 2.0 super resolution microscope.

Adult male Sprague-Dawley rats (200-250 grams) were randomly allocated into three groups: rats given a subcutaneous injection of phosphate buffered ssaline, rats treated with an intraperitoneal injection of 1×10^11^ vector genomes of adeno-associated virus serotype 9 encoding green fluorescent protein (AAV-GFP) one week after subcutaneous monocrotaline (MCT) (60 mg/kg) injection, and MCT rats treated with 1×10^11^ vector genomes of AAV9 encoding JPH2 (AAV-JPH2) one week post MCT injection. The cardiac-specific *TNT4* promoter directed expression for both viruses. RV mitochondrial enrichments were processed for quantitative proteomics using TMT10-plex labeling^5^. Lipidomic profiling of >1,100 lipid species in RV specimens was performed by Metabolon, Inc. Electron micrographs of RV mitochondria were collected at the University of Minnesota (UMN) Imaging Center. Blinded image analysis was performed by SZP or FP. Echocardiography and closed-chest pressure-volume loops^5^ defined RV function and pulmonary hypertension severity. Animal studies were approved by the UMN Institutional Animal Care and Use Committee. Statistical analyses were performed on GraphPad Prism 9.0 and MetaboAnalyst software (https://www.metaboanalyst.ca/).

Super resolution microscopy revealed discrete areas of JPH2 and MFN2 co-localization in RV cardiomyocytes (**Figure A**). Both JPH2 and MFN2 were detected in mitochondrial fractions, and coimmunoprecipitation experiments showed JPH2 and MFN2 interacted in RV extracts (**Figure A**). Pulldown studies demonstrated JPH2 directly bound MFN2, and the binding domain was mapped to the amino-terminal third of JPH2 (**Figure A**).

**Figure:**
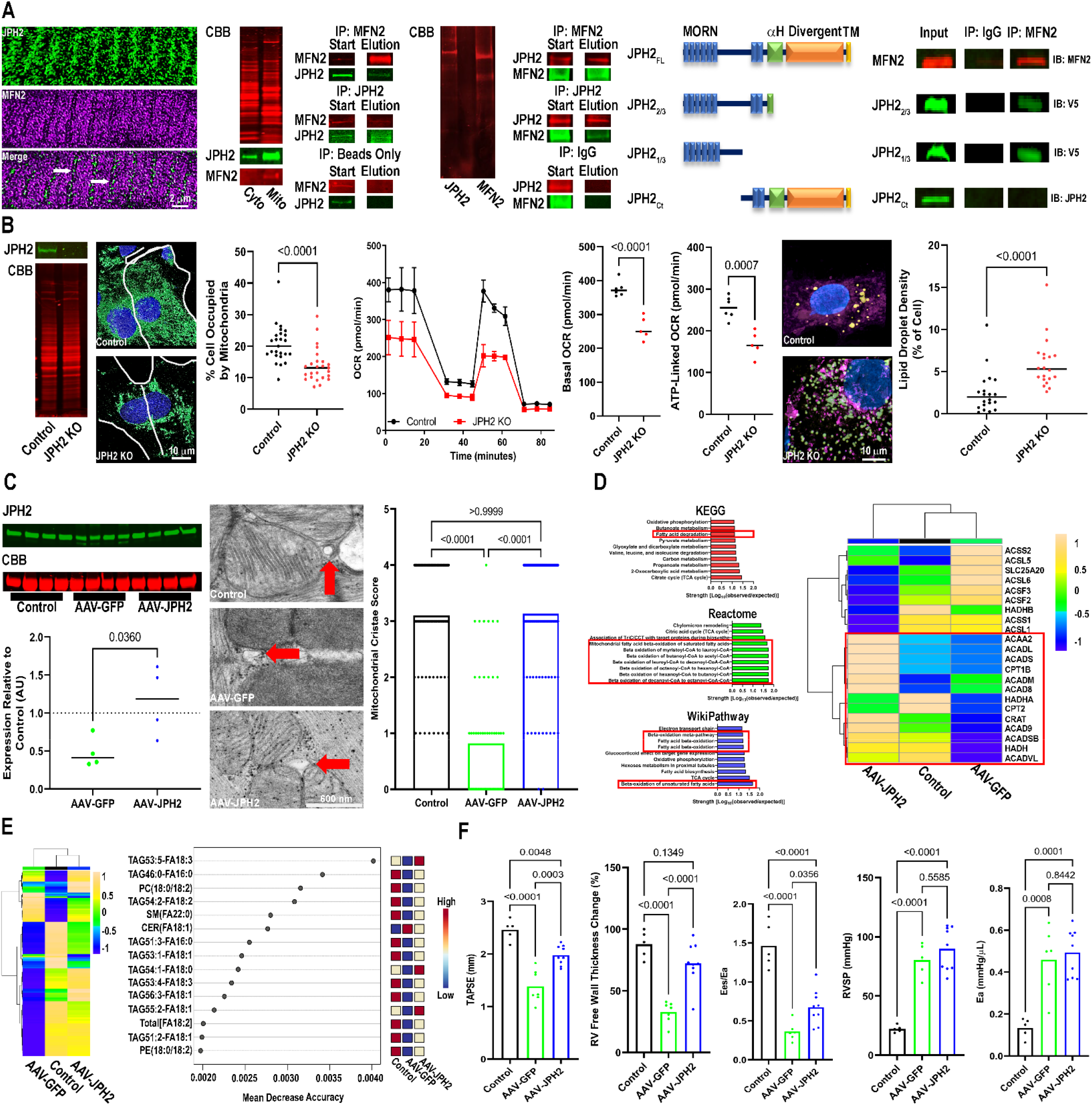
JPH2 binds MFN2 and regulates mitochondrial metabolism and RV function. (A) Confocal micrographs demonstrated JPH2 (green) and MFN2 (purple) co-localized in isolated RV cardiomyocytes. Coomassie brilliant blue (CBB) of SDS-PAGE of cytoplasmic and mitochondrial extracts. Western blot analysis of cellular fractionations showed both JPH2 and MFN2 were enriched in the mitochondrial fraction. Co-immunoprecipitation studies demonstrated a biochemical interaction between JPH2 and MFN2. Immunoblots of pulldown experiments of recombinantly expressed and purified JPH2 and MFN2 constructs revealed the two proteins directly bound each other, and the interaction site was localized to the amino-terminal third of JPH2. (B) Western blot analysis showed JPH2 expression was abolished in JPH2 KO iPSC-CM (above) and CBB gel showed equivalent loading (below). Representative confocal micrographs revealed a reduction in mitochondrial mass (*p*-values determined by unpaired *t*-test) in *JPH2* KO iPSC-CM. Seahorse tracings and quantification of reduced oxygen consumption rates (OCR) in *JPH2* KO iPSC-CM. Confocal micrographs of lipid droplets (left) and the area of cells occupied by lipid droplets in control and *JPH2* KO iPSC-CM. (C) AAV-JPH2 restored JPH2 protein abundance and improved peri-t-tubular (red arrow) mitochondrial cristae structure. (D) KEGG, Reactome, and WikiPathway analysis of differentially expressed proteins in mitochondrial enrichments. Red boxes highlight FAO related pathways. Hierachical cluster analysis demonstrated AAV-JPH2 increased mitochondrial FAO protein abundance in RV mitochondrial extracts. (E) Hierarchical cluster and Random Forest analyses suggested AAV-JPH2 treatment restructured RV lipid homeostasis. (F) AAV-JPH2 treatment enhanced RV function as assessed by echocardiography and pressure-volume loop analysis without significantly altering RV afterload.

Next, we evaluated the effects of JPH2 ablation on mitochondrial abundance and function in iPSC-CM. JPH2 knockout cells had reduced mitochondrial density and oxidative capacity (**Figure B**). Moreover, JPH2 deletion heightened lipid droplet accumulation following oleate, linoleate, and palmitate exposure (**Figure B**), which suggested fatty acid oxidation (FAO) was impaired.

Then, we determined how restoration of JPH2 expression via AAV9 impacted mitochondrial morphology, mitochondrial protein regulation, and lipid levels in the RV of MCT rats. AAV-JPH2 treatment restored JPH2 levels in the RV and rescued defects in peri-t-tubule mitochondrial cristae morphology (**Figure C**). KEGG, Reactome, and WikiPathway all identified FAO as an altered pathway in our mitochondrial proteomics analysis (**Figure D**). Hierarchical cluster analysis of fatty acid handling/metabolizing proteins showed AAV-GFP rats exhibited downregulation of multiple FAO enzymes when compared to control animals, but AAV-JPH2 increased levels of several FAO enzymes (**Figure D**). Consistent with a FAO enhancing effect, lipidomic profiling demonstrated AAV-JPH2 combatted dysregulation of several lipid species (**Figure E**).

Finally, we quantified the effects of AAV-JPH2 on RV function. Echocardiographic and closed-chest pressure-volume loop analyses revealed AAV-JPH2 augmented RV function without significantly altering pulmonary hypertension severity (**Figure F**).

In conclusion, we show JPH2 directly interacts with mitochondria via MFN2. Ablation of *JPH2* in human iPSC-CM suppresses mitochondrial biogenesis and oxidative capacity and impairs lipid metabolism. These results are recapitulated in MCT rats as AAV-JPH2 corrects mitochondrial morphological defects, increases expression of multiple FAO proteins, and restructures the RV lipidomic signature. These molecular changes lead to improvements in RV contractility. The newly described mitochondrial-enhancing effect of JPH2, in addition to its other important cellular functions, support the hypothesis that increasing JPH2 levels could combat RVD.

## Notes

### Competing Interest Statement

KWP served as a consultant to Edwards and receives grant funding from Bayer.

